# Natural LXRβ agonist stigmasterol confers protection against excitotoxicity after hypoxia- reoxygenation (H/R) injury via regulation of mitophagy in primary hippocampal neurons

**DOI:** 10.1101/707059

**Authors:** Md. Nazmul Haque, Md. Abdul Hannan, Raju Dash, Il Soo Moon

## Abstract

Ischemic brain injury represents insufficient oxygen supply to the brain and further damage occurs upon reoxygenation due to elevated intracellular levels excitatory neurotransmitter glutamate and subsequent production of reactive oxygen species (ROS) which has long been related to neuronal cell death of hippocampus brain region. Previously, using cell biological assay and transcriptomics analysis we reported that naturally occurring phytosterol Stigmasterol (ST) promotes brain development and function through the enhancement of neuronal cytoarchitectural complexity and functional maturation in rat hippocampal neurons by induction of immediate early genes (IEGs). In the present study we investigated the STs role in neuroprotection and found that ST also dose-dependently increased the neuronal viability in hypoxia reoxygenation (H/R) induced injury at hippocampal culture. ST, at an optimal concentration of 20 μM, significantly reduced the transport of vesicular glutamate (VGULT1), synaptic vesicle pool size, expression of GluN2B, rate of ROS formation (DCFDA) but restore mitochondrial membrane potential (JC1) and DNA fragmentation (H2AX) against H/R induced injury. More interestingly, ST also significantly induces the expression of autophagy marker protein LC3BII and the adaptor protein P62 but not HSC70 which indicates STs capability of induction of chaperon independent autophagy at H/R treated cultures. Furthermore densitometric analysis reveals ST also significantly increases PINK1 (PTEN induced protein kinase 1) expression therefore, indicates its role in mitophagy. In addition, molecular dynamic simulations study indicates that ST bind to LXRβ and forms hydrogen bonds with ASN239, GLU281, ARG319, THR316, SER278, ASN239 and SER278 residues at high occupancy with GLU281(20.21%) and ARG319 (21.04%,) residues, which is necessary for sterol binding to the LXRβ. Taken together these findings suggest that neuroprotective effect of ST might be associated with anti-excitatory and anti-oxidative actions on CNS neurons and could be a promising drug candidate for the treatment or prevention of ischemic stroke related neurological disorders.

## Introduction

The cerebral consequence of global ischemia is a serious neurologic complication in certain vulnerable brain areas, including the pyramidal neurons of hippocampal CA1 region in both patients and experimental animals [1]. Ischemia can induce an imbalance of neurotransmitter release such as glutamate and causes excitotoxicity due to the rapid and massive release and inhibition of reuptake as a result of energy failure [2]. The accumulation of glutamate over activates a plethora of downstream signalling pathways, many of which involve a surge in intracellular calcium influx that result in neuronal sensitization to excitotoxic cell death. Furthermore, reperfusion which is critical in the treatment of ischemic stroke may worsen brain tissue damage in excess of the injury caused by ischemia alone by massive production of reactive oxygen species (ROS). A line of evidence indicates that elevated levels of ROS directly induces subsequent damage to enzymes and other proteins, lipids, nucleic acids, the cytoskeleton, and cell membranes, resulting in decreased mitochondrial function and lipid per-oxidation in pathophysiology of cerebral ischemia and stroke [3]. Therefore, prevention of glutamate-mediated toxicity induced ROS production could be a central therapeutic target in cerebral ischemia; unfortunately, a long list of drugs with assorted mechanisms of action on glutamate receptors failed to show efficacy when assessed in clinical trials [4]. However, the contribution of glutamate-mediated toxicity to functional outcome after acute ischaemic stroke has not been forgotten and has been the subject of additional study for exploring agents that target both glutamate-mediated toxicity and oxidative stress may be an effective therapy for ischemic diseases. Over the past decade, the study of the biological activity of glutamate toxicity in the CNS has gained particular interest and sterols have recently been suggested to help in protection against glutamate-mediated toxicity and ROS [5–7].

Depending on the nature of the diet, most commonly available plant sterols are sitosterol (65%), campesterol (32%), and stigmasterol (3%) [8].Among them sterol like Stigmasterol (ST) which have high structural and functional similarity to cholesterol is found in many vegetables, nuts, legumes, seeds and herbs [9].It can easily cross blood brain barrier and serves as a precursor for the synthesis of androgens, corticoids-1, estrogens, progesterone and vitamin D3 [10, 11]. It is well established that ST reduce plasma cholesterol levels, ostensibly by inhibiting enterocytic cholesterol uptake through competition with dietary and biliary cholesterol for absorption [12]. Furthermore, it has protective effect on glutamatergic neurotoxicity and inhibitory effect on acetylcholinesterase (AChE) activity and also possesses enhancement of memory [13, 14]. In addition, ST also promotes antioxidant, antiinflammatory effects and protect against strychnine and leptazol induced convulsions [9]. In our previous study we have shown that ST promotes the early neuronal cyto architectural development and reelin mediated induction of IEGs via ERK/ CREB signaling and potentiate functional presynptic plasticity as well as enhances the formation of dendritic spine in hippocampal neuron [16, 17]. All these activities might prove beneficial for the healthy brain function. We therefore in the present study observed the STs effect on neuroprotection in primary rat hippocampal neuron after hypoxia/reoxygenation (H/R) injury similar to that of ischemia reperfusion (I/R).

## Materials and methods

### Primary culture of hippocampal neurons and compound treatment

Primary hippocampal cell cultures were prepared as previously described by (Haque and moon 2018). Briefly on the 13th day of pregnancy time pregnant Sprague-Dawley rats was brought and housed in the animal care room under controlled temperature and 12 hr light-dark cycle with access to enough food and water till the 19th day of pregnancy and sacrificed after euthanization with isofluorane, approved by the Institutional Animal Care and Use Committee of College of Medicine, Dongguk University. The rat embryos (gestational age: E18/19) were dissected and the fetuses were collected. After isolation of hippocampai form the fetuses brain it was dissociated to single cell using fine pasture pipette and the dissociated cells were seeded at of 3.0 × 10^4^ cells/cm^2^ for neuroprotective study. Treatment of Stigmasterol, (ST) or vehicle (DMSO, final concentration < 0.5%) was applied into the culture media prior to cell plating. Hypoxia/reoxygenation (H/R) treatment in neurons was carried out as followed by Mohibbullah et al. [17]. Briefly, after the indicated times, neurons on coverslips in the culture plates were transferred into a modular hypoxic incubation chamber (Modular Incubator Chamber MIC-101; Billups-Rothenberg Inc., Del Mar, CA, USA) under atmospheric conditions of 94% N2, 5% CO2, and 1% O2, and incubated at 37^0^C for 3□h. To re-oxygenate hypoxic cells, culture plates were returned to a normoxic incubator obtaining 95% air and 5% CO2 at 37°C, and incubated until 96h for future experiments. All the reagents used (unless otherwise stated) for primary cell cultures were purchased from Invitrogen (Carlsbad, CA, USA) unless or otherwise stated.

### Cell viability assay

Evaluation of neuronal viability was performed using trypan blue exclusion assay at the indicated time of cultures. The neurons on coverslips were stained with 0.4% trypan blue for 15 min at room temperature (RT) and then washed with Dulbecco’s phosphate buffered saline (D-PBS; Invitrogen). Trypan blue dye itself is a cell impermeable but only can enter the cells with compromised membrane, thus rendering the cells dark blue color in phase-contrast images. Viability is expressed as the percentage of trypan blue-impermeable cells (live neurons) and the results were normalized to trypan blue-stained controls (no H/R). To enhance the accuracy and reliability of the results of cell viability, additionally, MTT assay was performed. Briefly, MTT was added to the culture medium at a final concentration of 0.5 mg/ml and incubated at 37 °C for 4 h. Then the medium was discarded, and DMSO was added to solubilize the formazan reaction product with shaking for 5 min. The optical density (OD) was spectrophotometrically measured at 570 nm using a microplate reader (Molecular Device, Spectra Max 190, USA). Cell viability of the control group was defined as 100%. Other groups’ cell viabilities were expressed as a percentage of the normoxia (no H/R) control cells (100%).

### Measurement of mitochondrial membrane potential and intracellular ROS

The mitochondrial membrane potential (MMP) as an indicator for healthy mitochondria in neurons was quantitavely assessed with 5,5’,6,6’-tetrachloro-1,1’,3,3’-tetraethylbenzimidazolyl-carbocyanine iodide (JC-1) labeling (1 μg/mL; Molecular Probes). Neurons on coverslips were washed with pre-warmed fresh media, incubated with JC-1 for 30 min at 37°C, and further washed twice with pre-warmed fresh media. Fluorescence images were obtained at 488 nm (green) for JC-1 monomers and at 568 nm (red) for JC-1 aggregate. The intensity ratio of the red/green fluorescence of each neuron was used to measure the level of MMP using Image J (version 1.45; National Institutes of Health [NIH], Bethesda, MD, USA). Since dysfunction of MMP responsible for ROS production we therefore further determined the ROS production using the fluorescent probe DCFDA (Molecular Probes, Eugene, OR, USA) on DIV 13 with either control or ST. Briefly the cultures were exposed to 10 μM DCFDA for 30 min. under the dark condition in the incubator and washed with pre-warmed PBS and visualized under a fluorescence microscope and images were collected and quantified for ROS positive cells in FL-1 channel.

### FM1-43 staining

To evaluate the functional maturation of hippocampal neurons in culture, cycling synaptic vesicles were stained with the styryl dye N-(3-triethylammoniumpropyl)-4-(4-(dibutylamino) styryl) pyridium dibromide (FM1-43; Molecular Probes, Inc., Eugene, OR). For labeling of synaptic vesicles, ST or vehicle treated hippocampal neurons were incubated for 3 minutes in depolarizing extracellular solution (50 mM KCl, 54 mM NaCl, 2 mM CaCl2, 1 mM MgCl2, 20 mM HEPES; pH 7.3) containing 10 μM FM1-43 dye. The use of depolarizing solution induced vesicle exocitosis and facilitated the uptake of FM1-43. After incubation with FM1-43 dye, neurons were washed three times (5 minutes each) in extracellular solution (0 mM Ca^+2^/3 mM Mg^+2^) without FM1-43 dye to wash off excess dye. Typical fluorescence images of stained synaptic vesicles would be of FM1-43 entrapped scattered puncta.

### Immunocytochemistry and immunoblotting

Neurons on coverslips were rinsed briefly with D-PBS and fixed by a sequential paraformaldehyde/methanol fixation procedure (Moon et al. 2007) on DIV 13 that is after 96h of hypoxia. For immunostaining, The following antibodies were used: primary antibodies to tubulin α-subunit (mouse monoclonal 12G10, 1:1000 dilution; Developmental Studies Hybridoma Bank, University of Iowa, USA); anti-microtubule associated protein 2 (MAP2, 1:500, mouse monoclonal, Sigma, St.Louis, MO), GluN2B (1:1000, rabbit polyclonal) anti-phospho-H2AX antibody (1:500; Millipore, Billerica, MA, USA); anti-heat shock conjugated 70 (HSC70, 1:100, rabbit polyclonal, Proteintech, Chicago, IL), and secondary antibodies (Alexa Fluor 488-conjugated goat anti-mouse IgG [1:1,000] and Alexa Fluor 568-conjugated donkey anti-rabbit IgG [1:1,000], Molecular Probes). Fixed neurons were incubated with primary antibodies followed by secondary antibodies and mounted on slides as described (Moon et al. 2007). For immunoblotting hippocmapal cells (density 1.2×10^5^ cells/cm^2^ on DIV13) were harvested after H/R treatment and lysed in ice-cold RIPA buffer [50 mM Tris– HCl (pH 8.0), 150 mM NaCl, 1% (v/v) NP-40, 0.5% (w/v) sodium deoxycholate, 1% (w/v) sodium dodecylsulfate, and protease inhibitor cocktail (Thermo Scientific, Rockford, IL)]. After collection, protein concentrations were measured (Bradford, 1976) and equal amounts of protein were separated accordingly (Haque and Moon 2018b), then incubated with primary antibodies: rabbit polyclonal anti-LC3II, anti-P62 (1:2,000; rabbit polyclonal, Cell signaling, Denvers, MA), anti-HSC70(1:1500, rabbit polyclonal, Proteintech, Chicago, IL), and anti-actin (JLA20; 1:1,000, mouse monoclonal, Developmental Studies Hybridoma Bank, University of Iowa, IO). After rinsing with TTBS (0.05% Tween-20 in TBS), membranes were incubated with horse radish peroxidase-conjugated secondary antibodies (1:1,000; anti-mouse or -rabbit IgG; Amersham Biosceinces), and blots were detected using an ECL detection kit (Invitrogen, Waltham, MA).

### Molecular dynamics simulation

The crystal of human LXR Ligand Binding Domain domain was retrieved from the protein data bank (PDB ID: 1P8D) and prepared by using the bond orders, hydrogen and charges. The structure was refined by removing water molecules and optimizing the protein at neutral pH. Readjustment of some thiol and hydroxyl groups, amide groups of asparagines, glutamines and imidazole ring of histidines, protonation states of histidines, aspartic acids and glutamic acids were done. Minimization was done by applying OPLS 3 force field by adjusting maximum heavy atom RMSD to 0.30 Å. In order to perform Glide XP docking, the three dimensional coordinates of Stigmasterol was downloaded from PubChem databases and prepared by using Ligand preparation wizard of Maestro 11.1 [19] with an OPLS 3 force field. Their ionization states were generated at pH 7.0±2.0 using Epik 2.2 in Schrödinger Suite 2017-1. The active site of the protein was fixed for docking simulation by generating a grid box at the reference ligand binding of the protein. Grid generation parameters were kept at default with a box size of 18 Å × 18 Å × 18 Å, and the OPLS 3 force field utilized for post minimization. The charge cutoff and vander Waals scaling factor were set to 0.25 and 1.00 respectively [20]. We carried out extra precision (XP) flexible docking by Glide module of Schrödinger-Maestro v9.4 [21, 22]. Here, all ligands were treated flexibly with considering the partial charge and ven der Waals factor of 0.15 and 0.80 respectively. Minimization was performed to the docked complex after docking using the OPLS_2005 force field, where each ligand. Additionally, Prime MM-GBSA approach was carried out for calculating binding free energy which combines OPLSAA molecular mechanics energies (EMM), an SGB solvation model for polar solvation (GSGB), and a non-polar solvation term (GNP) composed of the non-polar solvent accessible surface area and van der Waals interactions [23]. The total free energy of binding: ΔGbind = Gcomplex − (Gprotein + Gligand), where G = EMM + GSGB + GNP After that, the docked complex and ligand free protein were subjected for molecular dynamics simulations using YASARA Dynamic software. Prior to the simulation, all structures were cleaned and subjected for the optimization of hydrogen bonding network. For each simulation system, a cubic simulation cell with periodic boundary condition was generated and then all atoms were parameterized with the AMBER14 [24] force field. The transferable intermolecular potential3 points (TIP3P) water model was used to make solvation system, and the density was maintained to 0.997 gm/L. Using the simulated annealing method, Initial energy minimization process of the each simulation system was done by using steepest gradient approach for 5000 cycles. Molecular dynamics simulations were done by using the PME methods to describe long-range electrostatic interactions at a cut off distance of 8 Å at physiological conditions (298 K, pH 7.4, 0.9% NaCl) [25]. A multiple time step algorithm along with a simulation time step interval of 2.50 fs was chosen [26]. At a constant pressure and Berendsen thermostat, molecular dynamics simulations were performed for 100 ns long, and MD trajectories were saved every 25 ps for further analysis.

### Image acquisition

A Leica Research Microscope DM IRE2 equipped with I3 S, N2.1S, and Y5 filter systems (Leica Microsystems AG, Wetzlar, Germany) and a high-resolution CoolSNAP™ CCD camera (Photometrics Inc., Tucson, AZ, USA) was used for phase-contrast and epifluorescence microscopy, through Leica FW4000 software. The digital images were processed using Adobe Photoshop 7.0 software. Quantitative analysis of staining intensity of FM1-43 labeled recycling synaptic vesicles, and VgulT1, GlunN2b Puncta per 50 μm length of dendrite was meansured respectively by fluorescence intensity of individual stained punctae with Image J (version 1.45) (National Institute of Health, Bethesda, MD, USA) and puncta analyzer [27] software.

### Data analysis

All data are expressed as the mean ± SEM of at least three independent experiments. Statistical comparisons were made by one-way analysis of variance (ANOVA) with post hoc Duncan multiple comparisons and Bonferroni correction (SPSS software, version 16.0). Predetermined p values ≤ 0.05 were considered statistically significant.

## Results

### Optimization of STs Concentration for neuroprotection

To investigate the effects of ST on cell viability, a trypan blue exclusion assay was carried out with primary hippocampal cells. Incubation with ST at various concentrations up to 225 μM changed the cell viability compared to vehicle only (EtoH)-treated cells (Fig. 1). ST at concentration range of 2.50-150 μM significantly increased cell viability and with an optimum concentration of 20 μM. In addition MTT [3-(4, 5-dimethylthiazol-2yl)-2,5-diphenyltetrazolium bromide] assay also shows the similar trend of viability to that of trypan blue. We therefore used dose of 20 μM of ST in all further experiments.

**Figure 1.**
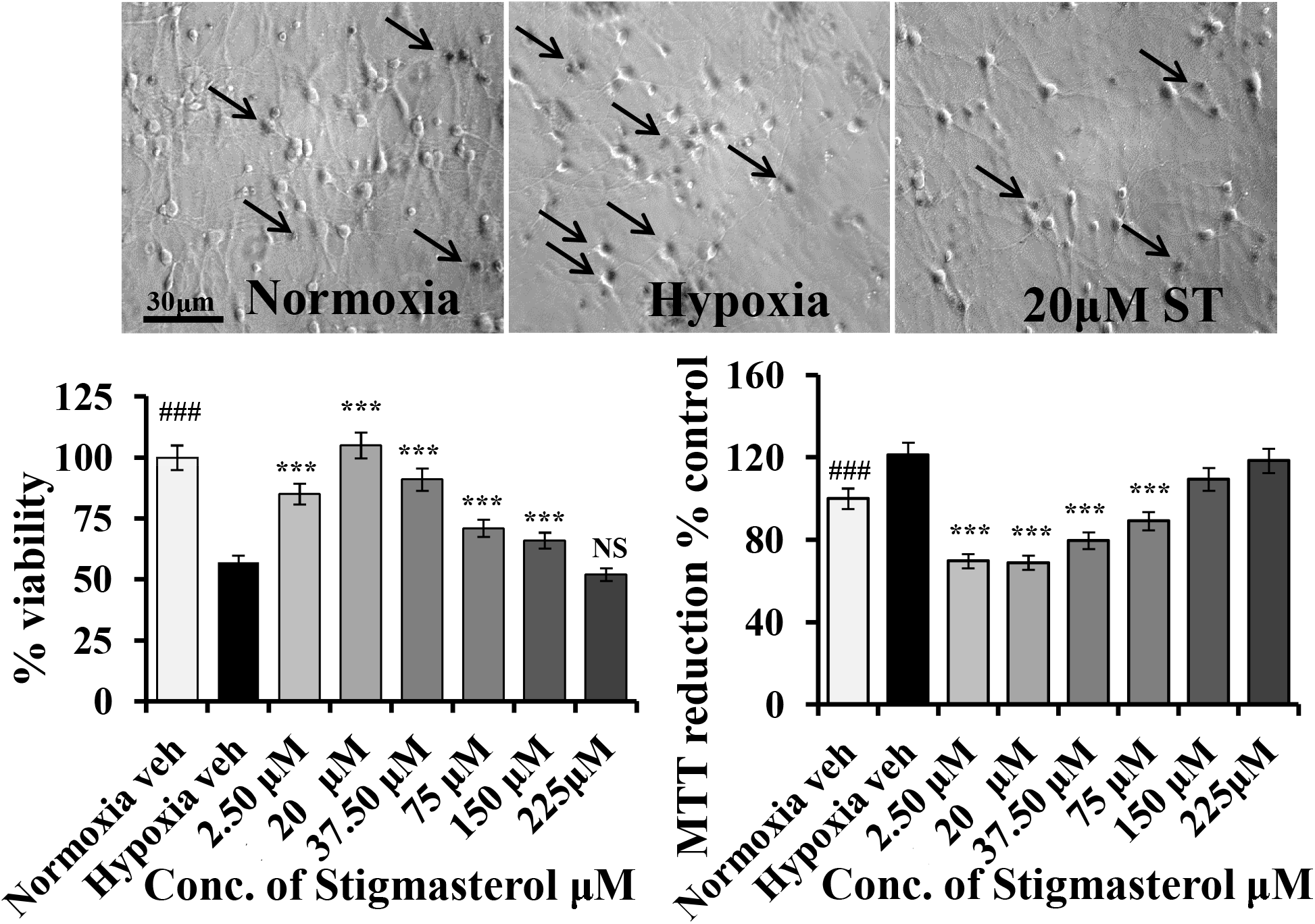
Effect of ST on neuronal viability after H/R and normal cultures. Embryonic (E) 19-d rat hippocampal cells were cultured on poly-DL-lysine-coated coverslips in serum-free neurobasal media supplemented with B27, and plated at 3.0 × 10^4^ cells/cm^2^. On DIV 9, cultures were exposed to hypoxia (3 h) and subsequently to normoxia (96 h). The viability was determined by trypan blue exclusion and MTT assays in both hypoxic with or without ST treated cultures, and normalized to vehicle-treated normoxic controls. Bars represent means ± SE (n = 3; ~800–1,000 neurons per group for staining). ^#^*p* < 0.05, ^###^*p* < 0.001 compared with the normoxia control; \**p* < 0.05, \*\*\**p* < 0.001 compared with the hypoxia control (ANOVA).

### ST decreases synaptic vesicle pool size and extrasynaptic GULN2B receptor expression after H/R

Hypoxia/reoxygenation induces hyperexcitability of neurons by glutametergic signaling i.e. AMPA, NMDA activation [28–30]. Since exocytotic release of glutamate depends upon loading of the neurotransmitter into synaptic vesicles by vesicular glutamate transporters, VGLUT1, which also determines the quantal size via controlling the kinetics of synaptic vesicle recycling and the probability of release of neurotransmitter [31, 32]. We therefore, in the present study analyzed the expression of VGLUT1 by immunostaining and quantal size by FM1-43 live stain at DIV13 and found that after H/R [(−) ST H/R], there was an increase in VULT1 puncta expression by 55% compared to normoxia control [(−) ST no H/R, P<.001] which was successfully attenuated by ST [(+) ST H/R, P<.001] treatment (Fig. 2A). Moreover, glutamate mediates the expansion of release mercenary in presynaptic terminal, quantification of the signal intensity of FM1-43 labeled cycling synaptic vesicles from [(−) ST no H/R], [(−) ST H/R] and [(+) ST H/R] cultures (Fig. 2B) it appear that the mean fluorescence intensity of the FM1-43 dye accumulations in reserve vesicle pool was significantly higher (about 1.15 folds, P<.001) after H/R [(−) ST H/R] compare to [(−) ST no H/R] indicating that [(−) ST H/R] treated neurons are more active in synaptic transmission which is compensated by ST treatment [(+) ST H/R, P<.001]. Furthermore, evidences suggest that glutamate spillover and activation of extrasynaptic receptors occurs when large glutamate amounts are released from synaptic endings [33].We therefore, measured the expression level of extrasynaptic glutamate receptor GULN2B using both immunocytochemistry and immunoblotting and found that under ST treated hypoxic condition there slight change in GULN2B puncta expression compare to normoxic condition (around 7%) but under H/R condition increases in GULN2B expression significantly higher (by 30% vs. normoxic control, P<.001 and by 21% vs. ST treated hypoxic condition, P<.001 respectively) and blotting also shows increase in densitometric expression of GULN2B after H/R (Fig 2C).

**Figure 2.**
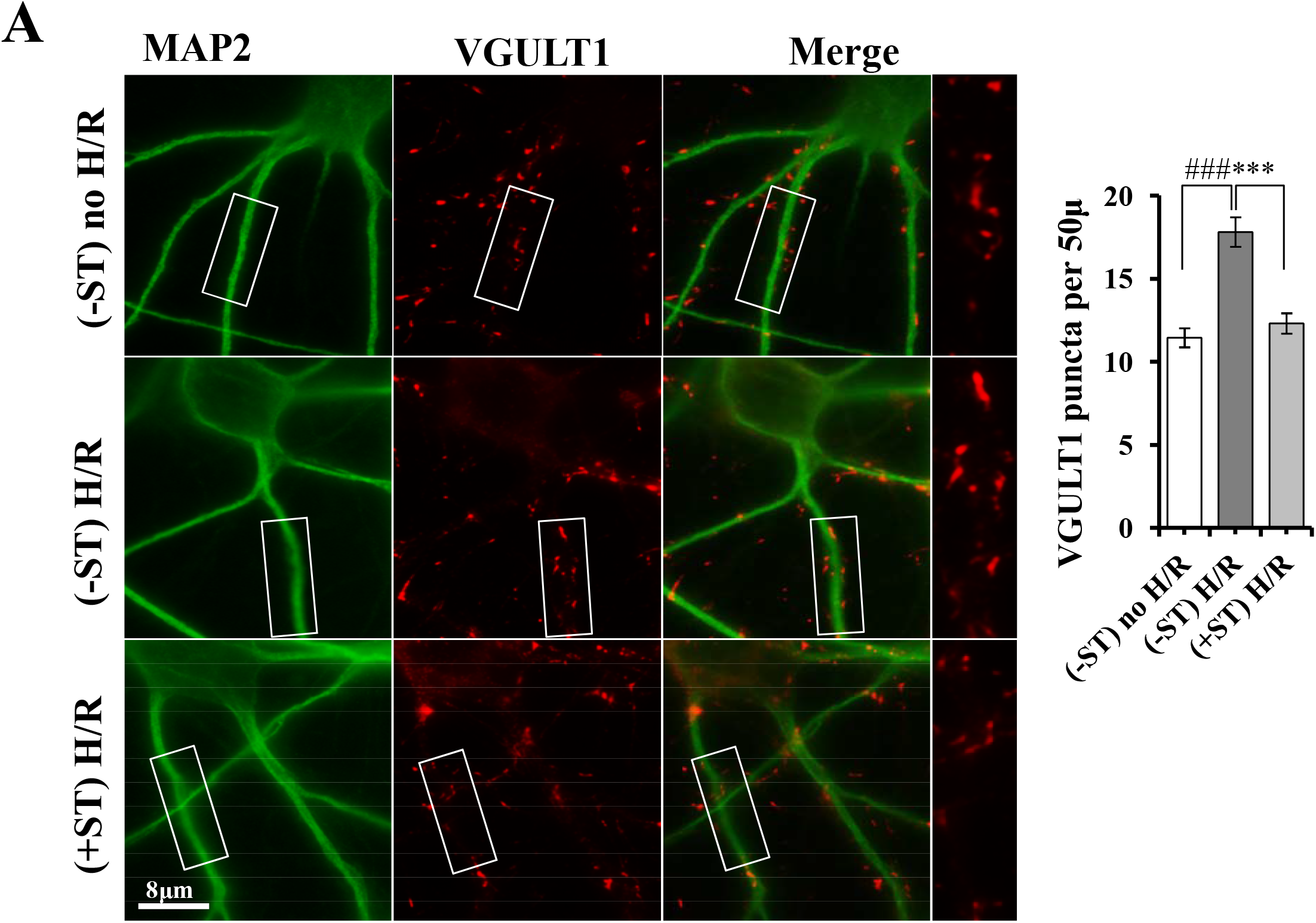
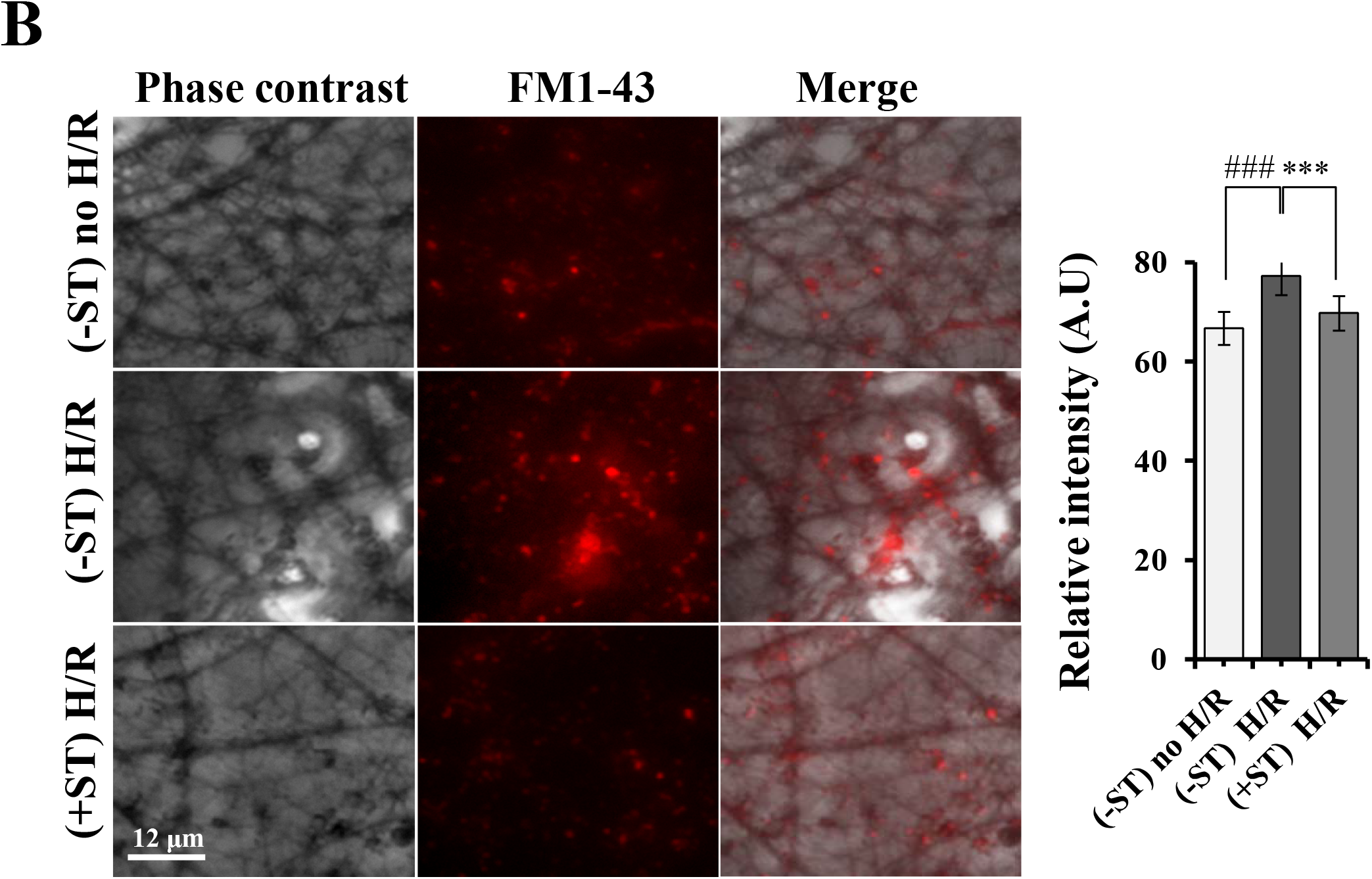
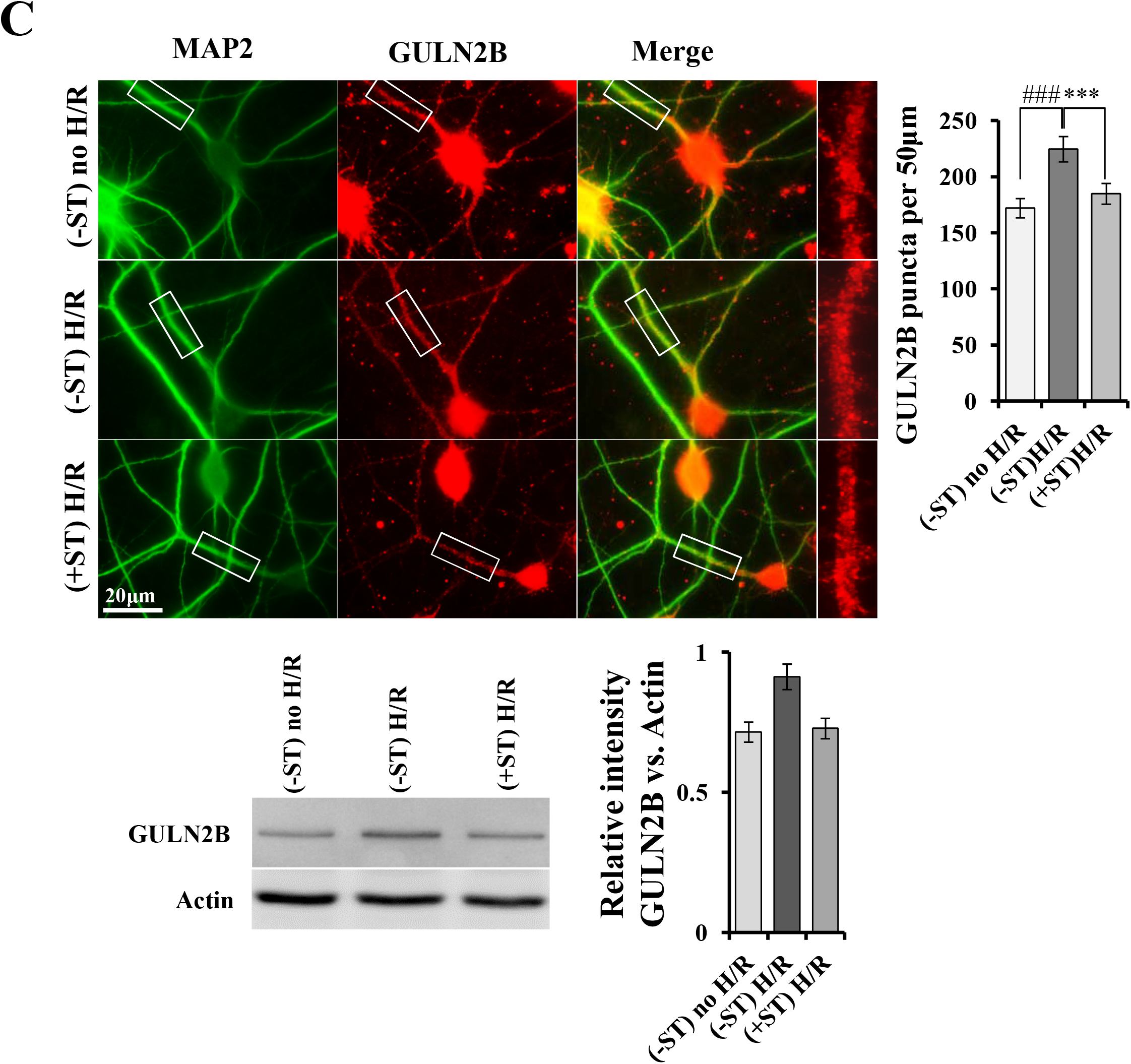
Effect of ST on pre-and post synaptic neurotransmission (glutamate) after H/R. Vehicle or ST (20μM) was added to the hippocampal cultures. On DIV 9, the cultures were exposed to hypoxia, then to normoxia for 94 h and fixed, double-immunostained for anti MAP2 (green)/VGLUT1 presynaptic (red), and MAP2 (green)/GluN2B postynaptic (red) antibodies. (A) Representative immunofluorescent images of VGLUT1 punctae of normoxic control [(−) ST no H/R)], hypoxia control [(−) ST H/R)] and ST-treated [(+) ST H/R) cultures and their statistics. (B) FM1-43 stained synaptic vesicle accumulations in dendritic areas and statistics of mean intensities (arbitrary unit, a.u.) of FM1-43 labeled punctae, (n = 3; 30 microscophic field per group). (C) Immunofluorescent and blotting expression of GluN2B and their statistics. Bars represent means ± SE (n = 3; 30 neurons per group). ^#^*p* < 0.05, ^###^*p* < 0.001 compared with the normoxia [(−) ST no H/R)] control; \**p* < 0.01; \*\**p* < 0.05 compared the hypoxia [(−) ST H/R)] control (ANOVA).

### ST Increases MMP and decreases ROS production H/R

It is well known that glutamate can stimulate ROS formation or vise versa which mediates via NMDA receptors after H/R through mitochondrial dysfunction [34–36]. Since ST is reported to increase glutathione content [37] which provide an indication of antioxidant defense mechanism and may protect mitochondrial function. We, therefore, observe the effect of ST on MMP maintenance after H/R. Hippocampal cultures were exposed to hypoxic conditions on DIV 9 and MMP levels were measured using JC-1 (a fluorescent probe) after 96 h of reoxygenation. Before H/R, cells grown without ST [(−) ST no H/R] showed a mixture of green (representing JC-monomer i.e., low MMP) and red fluorescing (representing JC-aggregate i.e., high MMP) mitochondria (Fig. 3A). When (−) ST H/R cultures were exposed to H/R, they showed mostly green fluorescence [(−) ST H/R], whereas ST-treated cultures [(+) ST H/R] showed mostly red fluorescence. The overall field intensities were measured using ImageJ software, and the intensity ratios of red to green fluorescence are shown in Figure 3A. H/R [(−) ST H/R, (3 h/96 h)] decreased the red/green ratio significantly (P<.05) to 1.14 − 0.09 (Fig. 3A), from 1.28 − 0.05 in normoxia control cultures [Fig. 8B, (−) ST no H/R]. In contrast, the red/green ratio was 1.27 – 0.06 in (+) ST H/R cultures under the same conditions, which represented a significant (P<.05) increase of MMP. These results indicate that ST inhibits MMP dissipation from mitochondria after H/R shock. Furthermore, as ST protects MMP therefore we address further question about STs role in ROS production under the same condition. As shown in Figure 3b, numbers of ROS-positive cells were increased by increasing hypoxia in both control [H/R and no H/R of ST (−)] and treatment with ST [(+) ST H/R] significantly (P < .001) decreased the numbers of ROS-positive cells by 10.12% compare to vehicle culture [(−) ST H/R] (Fig. 3B).

**Figure 3.**
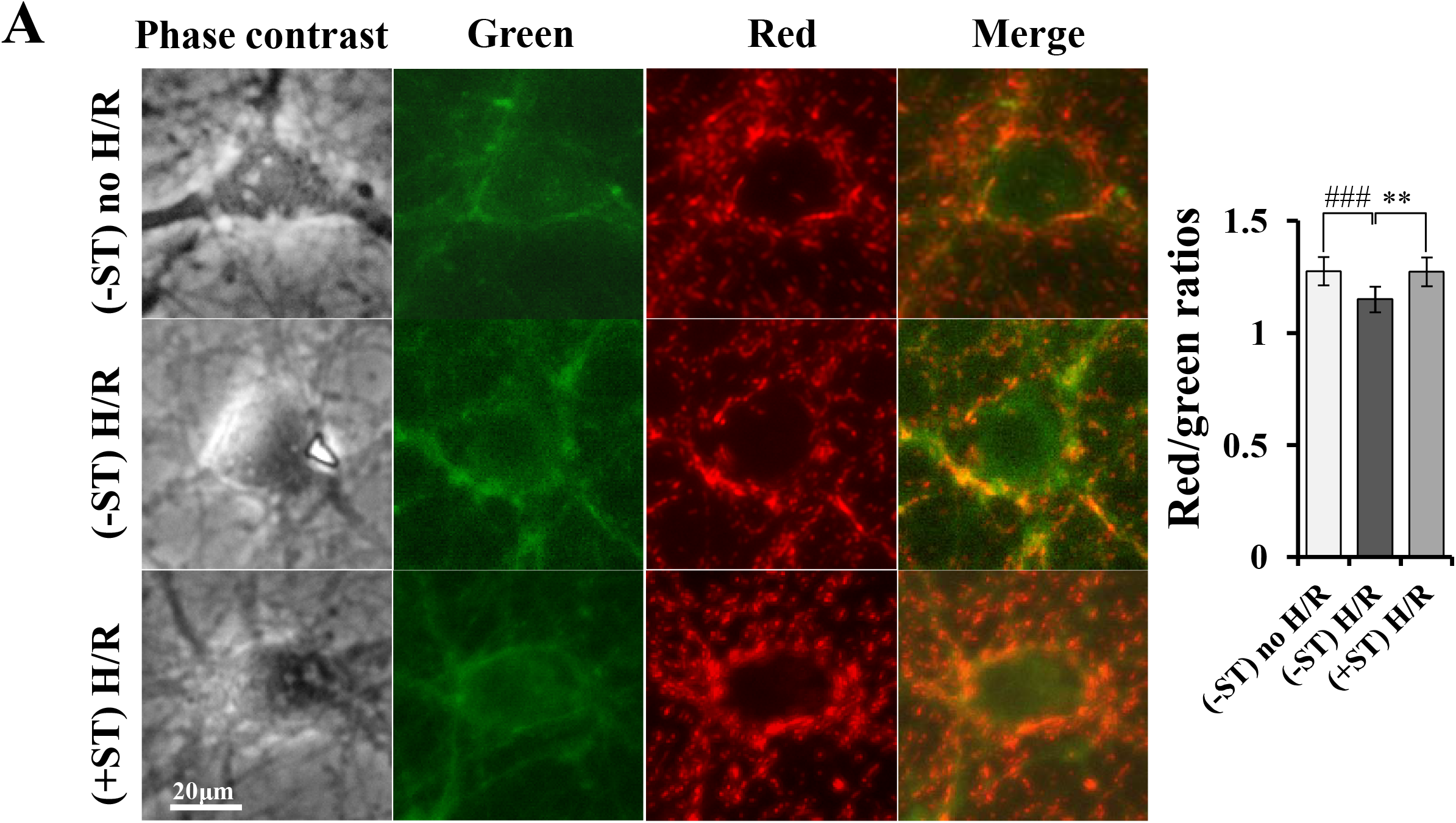
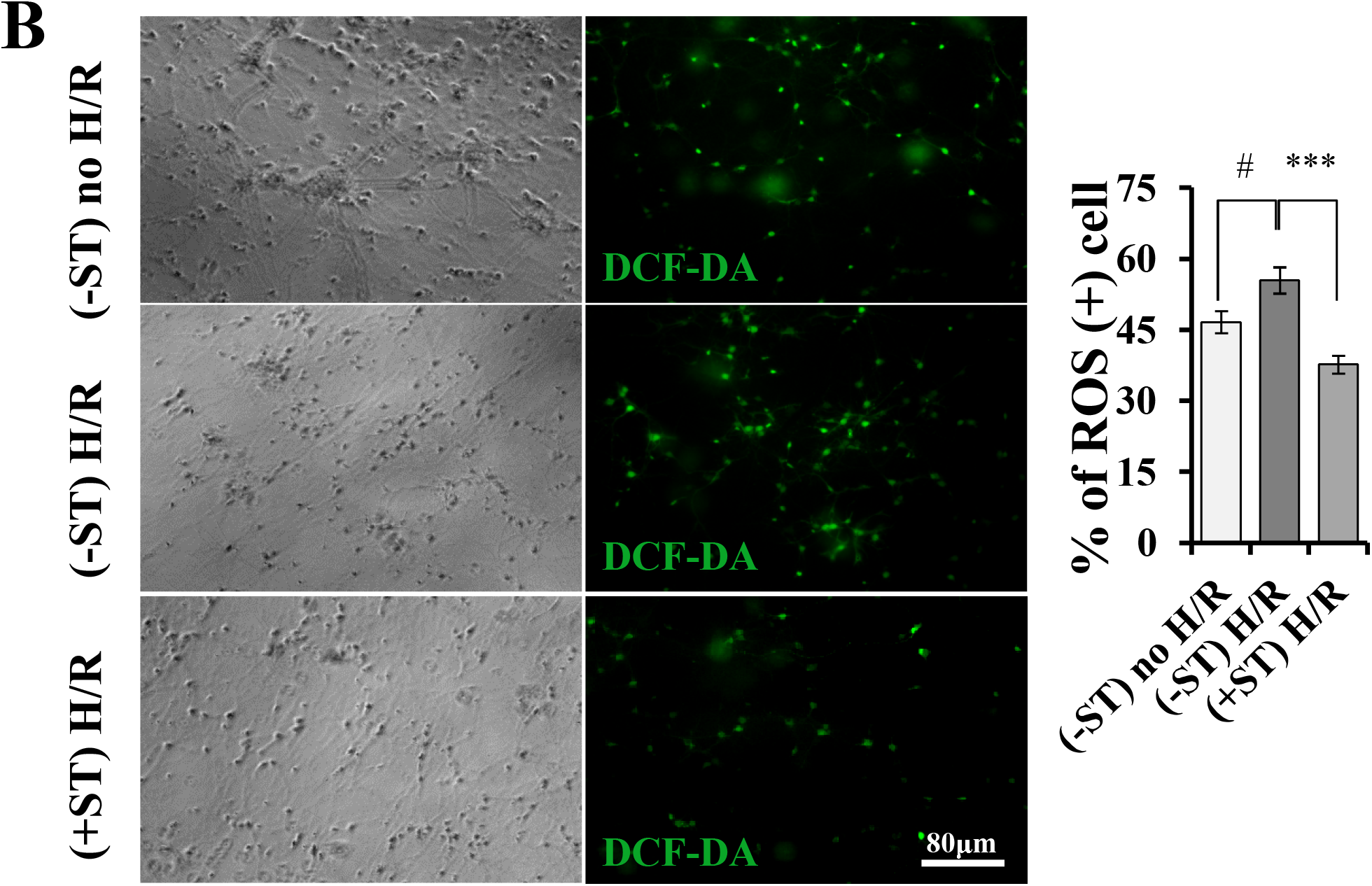
Effect of ST on MMP and ROS production after H/R. Vehicle or ST was added to the hippocampal cultures. On DIV 9, the cultures were exposed to hypoxia and then returned to normoxia. The MMP levels in cultured cells were assessed by staining with the fluorescent probe JC-1. (A) Representative images of JC-1 staining and statistics of red/green intensity ratios. Scale bar = 8 μm. Bars represent means ± SE (n = 3; 40 neurons per group). ROS levels were assessed by staining with the fluorescent probe DCF-DA. (B) Typical DCF-DA staining statistics of percentage of ROS (+) cells. Bars represent means ± SE (n = 3; ~1,000– 1,200 neurons per group). ^#^*p* < 0.05 compared with the normoxia control; \**p* < 0.01; \*\**p* < 0.05 compared with hypoxia control (ANOVA).

### ST Induces autophagy after H/R

Autophagy a pro-survival mechanism of cell and once induced by DNA injury, makes a crucial contribution in regulating cell fate [37, 38]. For example, many lines of evidence argue that autophagy can sustain the energy demand requires for DNA repair processes to delay apoptotic cell death upon DNA damage,[39, 40] by degrading toxic aggregates, elimination of putative sources of ROS, reduces their levels and, indirectly, decreases DNA damage accumulation[41–43].We therefore in the present study assed the STs capability for the induction of autophagy after H/R and found that ST [(+) ST H/R], increases the densitometric expression of autophagy induction protein LC3IIB (P<.05) western blot band (Fig. 4A) which is statistically significant compare to normoxia [(−) ST no H/R] (P<.05), as well as hypoxic condition [(−) ST H/R] (P<.05). Furthermore, ST also insignificantly enhances the expression P-62 (Fig. 4A). To dig deeper whether this autophagosome formation is chaperon dependent or not (Fig. 4A and 4B) we observed the expression of heat shock proteins (Hsp) family protein heat-shock cognate 70 (Hsc70), i.e. one of the critical components of the cell stress response and catalyze protein folding and trafficking and plays a central role in the protection of cells from damage via induction of signaling of cell survival and prevent apoptosis [44, 45]. In the present study we found that ST treatment didn’t influence the expression of HSC70. Since ST act similar to LXR agonist [46] and LXR agonist induces the expression of PINK1 [47] that is responsible for mitophagy, we therefore farther, analyzed the expression of PINK1 after H/R and found that ST significantly (P<.05) increases the densitometric expression of PINK1 [(+) ST H/R, (P<.05)] compare to vehicle control [(−) ST H/R] (Fig. 4A).

**Figure 4.**
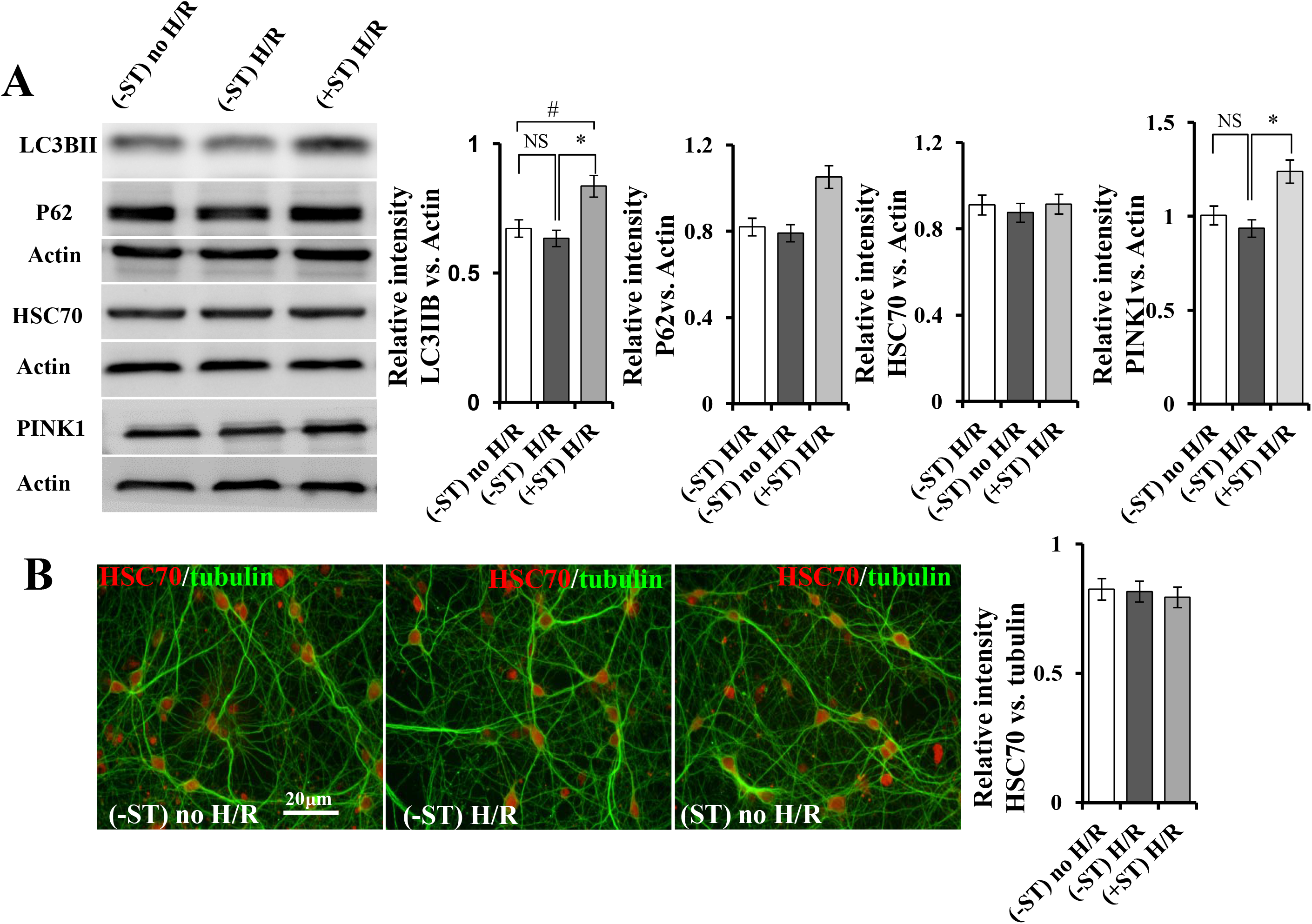
Immunoblot analysis on the effect of ST on LC3II and PINK1 expression after H/R. Vehicle or ST was added to the hippocampal cultures that is plated at 1.2 × 10^5^ cells/cm^2^ density and proteins were isolated and immunoblotted following hypoxia on DIV9 and then returned to normoxia for 96 h. (A) Representative immunoblot bands showing LC3II, P62, HSC70, PINK1 and Actin expressions along with their band intensities which was measured using Image J software and normalized to Actin. (B) Representative immunoflueroscence images of Tubulin (green)/HSC70 (red) staining and the statistics of relative intensity. Bars represent means ± SE (n = 3; 30 microscophic field per group for immunoflurescence images). ^###^*p* < 0.001 compared with the normoxia control; \**p* < 0.01; \*\**p* < 0.05 compared with hypoxia control (ANOVA).

### Protect DNA fragmentation

Dysfunction of chromatin due to ROS-mediated single and double-strand DNA break leads to cell death which is associated with apoptosis and necrosis. In view of that, hippocampal cultures were exposed to hypoxic conditions for 3 h on DIV 9, and immunostained after 96 h of reoxygenation with anti-phospho γ-H2AX antibody [48] a well-known marker of the DSB that mediate accumulation of various signaling and repair proteins to the damaged sites (Fig. 5). In controls [(−) ST H/R]), numbers of phospho-H2AX immunostained puncta per nucleus significantly (P < .001) increased from 10.5 − 4.9 [(−) ST no H/R]), to 23.55 − 7.9 after 96 h of reoxygenation. Pretreatment with ST (20 μM) significantly (P < .001) decreased numbers of puncta to 11.75 − 5.1 after 96 h of reoxygenation, respectively (Fig. 5).

**Figure 5.**
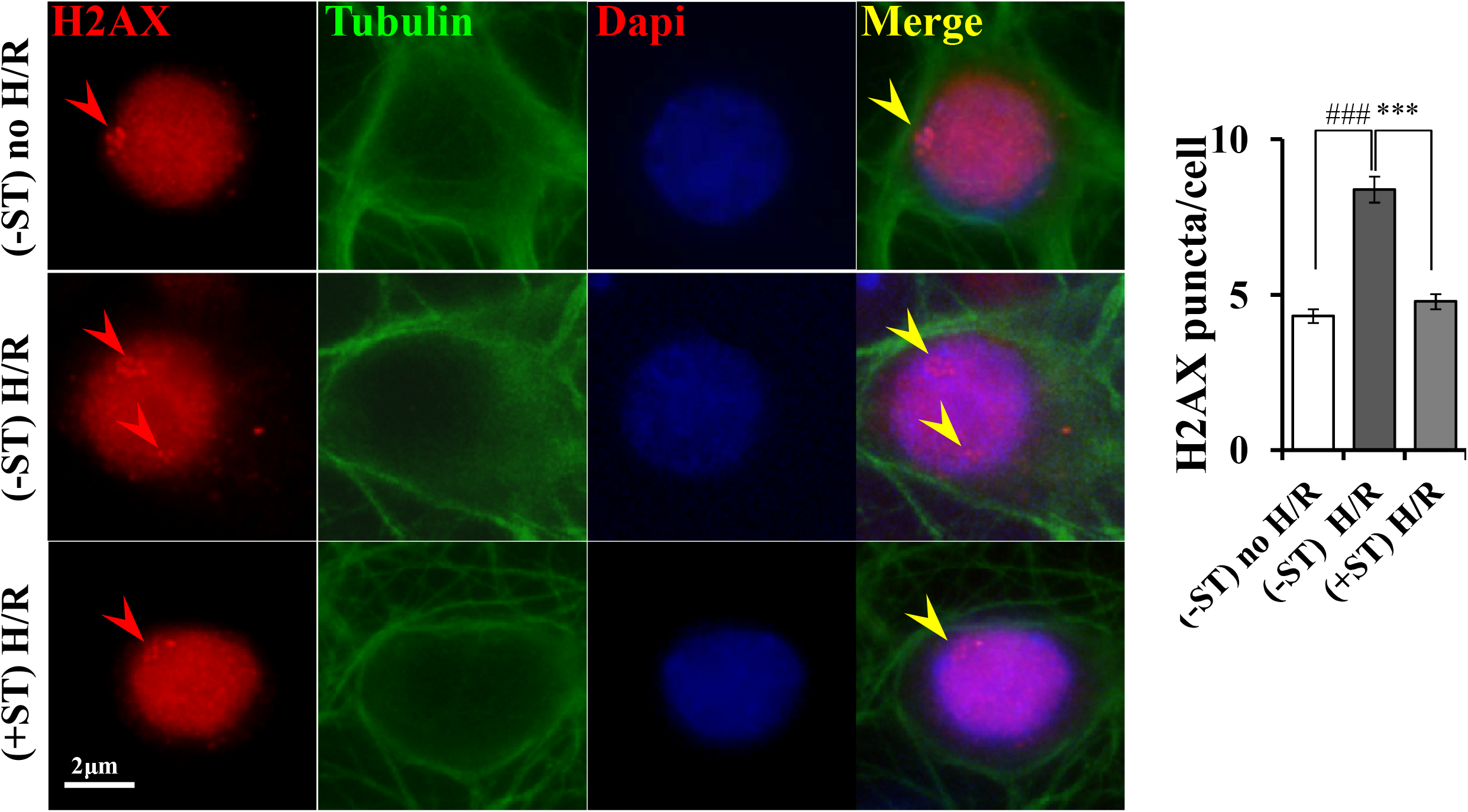
Effect of ST on DNA fragmentation after H/R. Hippocampal cells were cultured under the conditions described in Fig. 1. Cultures were exposed to hypoxia on DIV 9 and then to normoxia. Cells were immunostained with the anti-phospho-H2AX antibody. Representative immunofluorescent images. Nuclei are enlarged to show individual puncta (arrowheads) and statistics of numbers of punctae per nucleus. Bars represent means ± SE (n = 30 nuclei). ^###^*p* < 0.001 compared with the normoxia control; \**p* < 0.01; \*\**p* < 0.05.

### LXRβ agonistic conformation of ST by molecular dynamic simulation study

In order to reveal the binding interaction and orientation ST to the LBD domain of LXRβ, molecular docking simulation was performed by Glide XP docking. The docking study initially suggested that the ST fit well to the binding site and showed similar binding orientation to co-crystal ligand, desmosterol, a known agonist of LXRβ [49] (Fig. 6a). In detailed molecular interaction analysis revealed that ST formed several non-bonded interactions with the amino acids around the ligand binding domain, including two hydrogen bonds and several other hydrophobic interactions. ST was found to maintain hydrogen bonding with the ASN239 and PHE329 via its hydroxyl group in position C-3 of the steroid skeleton. Furthermore, the residues, PHE243, PHE271, THR272, LEU274, ILE309, MET312, PHE329, LEU345 and TRP 457 formed van der waals interaction with the ST by pi-alkyl bonding. The binding energy of ST to LXRβ was calculated and compared with the co-crystal ligand (Table S2). Interestingly, ST showed better binding energy than the native agonist with binding energy of −89.76 kcal/mol, while desmosterol, the cocrystal ligand, showed binding energy of −86.35 kcal/mol. The agonistic conformation of ST was further validated by 100 ns molecular dynamics simulation. In the simulation, the ligand free conformation of LXRβ was also included to check the conformational changes in LXRβ induced by ST binding. RMSD and RMSF value of the protein backbone (Cα atoms) of the two systems were calculated individually and plotted by comparing with the initial trajectory of protein structure. Both analyses revealed that binding of ST remained stable over the simulation, and achieved equilibrium after 50ns of total simulation time (Fig. 6b). Moreover, no significant local changes were found after binding of the ligand in the simulations, which signifies that ST formed stable binding to the receptor (Fig. 6c). In addition to that, the intermolecular interactions, in terms of hydrogen bonding was further calculated and rendered in Fig. 6d. The results showed that the total number of hydrogen bond changes from the starting simulation time, however, maintained over the simulation. Although molecular docking revealed the hydrogen bonding by ASN239 and PHE329 residue, only ASN239 showed interaction in the simulation, including others are GLU281, ARG319, THR316, SER278, ASN239 and SER278. The high h-bond occupancy was found for GLU281 and ARG319 residues, that is, 20.21% and 21.04%, respectively. However, ASN239 showed only 7.60% of interaction in the total simulation times. Since the hydrogen bonding interactions with GLU281 and ARG319 are necessary for sterol binding to the LXRβ, these results concluded the agonistic conformation of ST in LXRβ activation [49].

**Figure 6.**
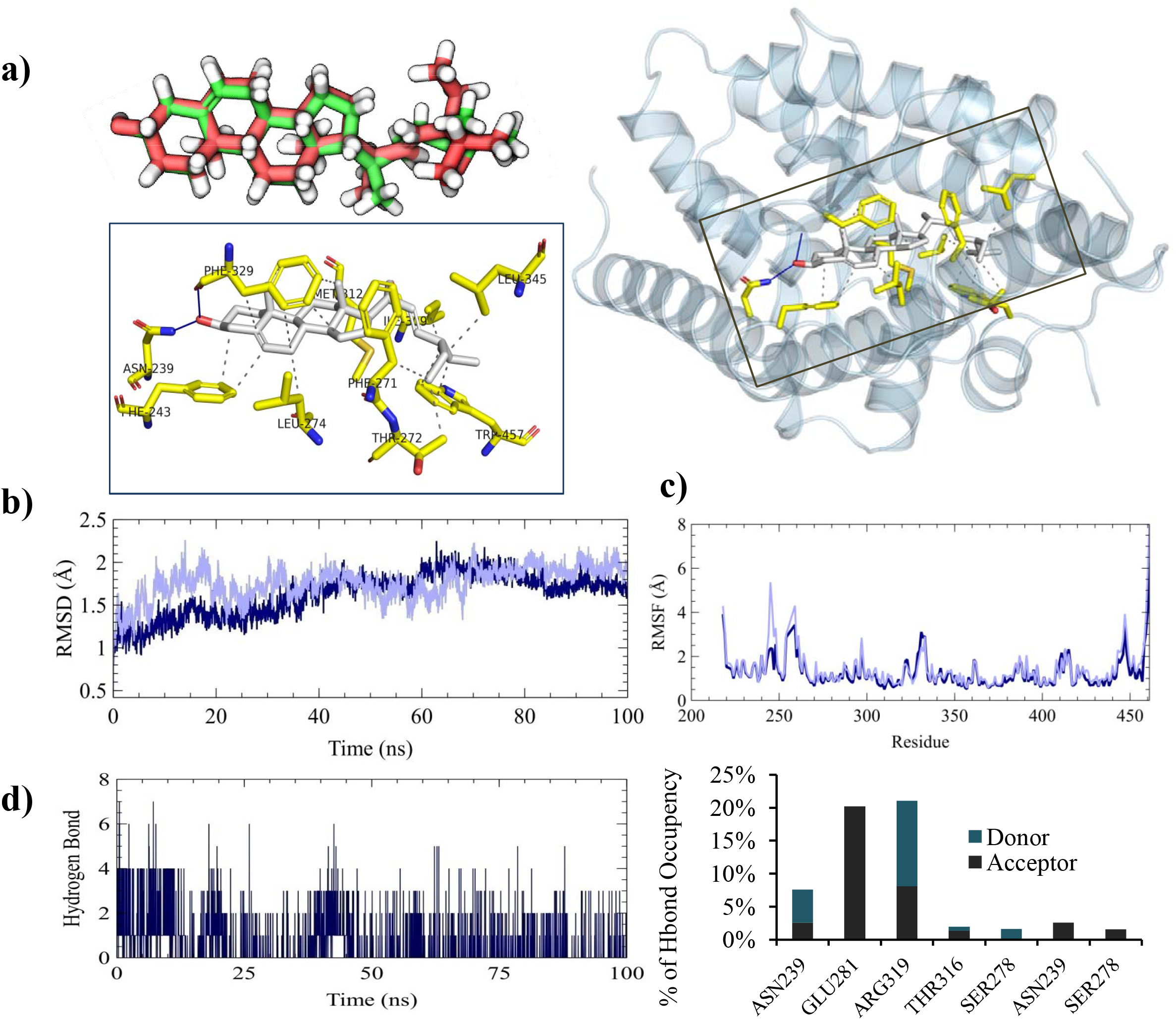

## Discussion

Brain injury by transient global brain ischemia (cardiac arrest) and focal brain ischemia (ischemia stroke) is the leading cause of serious and long-term disability worldwide. The hippocampus is responsible for many central nervous system functions including cognition, learning, and memory, but it is also one of the most vulnerable brain regions as regards to various neurological insults including hypoxia–ischemia [50]. Therefore protection against exacerbation of hypoxic injury after restoration of oxygenation (reoxygenation) could be one of the most important mechanisms for halting cellular damage in ischemia-reperfusion injury. Based on these considerations, we have explored the neuroprotective effects of naturally occurring phytosterol ST to in vitro H/R model that is mimetic to brain ischemia in the present study.

Glutamate, one of the major excitatory neurotransmitters in the mammalian central nervous system has direct effects on neuronal activity regarding brain function and toxic insults by glutamate in neuronal cell culture after H/R mimic a key component of ischemic brain injury. In addition microdialysis studies have shown that there is a several fold increase in extracellular glutamate during global ischemia, and could be a powerful neurotoxin and capable of killing neurons [51, 52]. In the present study we found that after H/R [(−) ST H/R)] in vehicle culture it surges glutamate like ischemic injury compare to that of no H/R [(−) ST no H/R)] and ST treatment [(+) ST H/R) attunes this excitotoxicity as evidenced by the increases in expression of Vgult1, ready to release synaptic vesicle pool size as well as extra synaptic glutamate receptor Glun2B which are similar to the previous findings as ST protect glutamate toxicity [5]. Moreover, under H/R conditions, the prolonged activation of extrasynaptic glutamate receptors can, causes a strong decrease in mitochondrial membrane potential, which can be considered to be an early marker for glutamte-induced neuronal damage, ROS generation [53, 54, 55] and cell death [55]. In this study we have found that ST treatment counterbalances the mitochondrial membrane potentiality and decreases the number of ROS positive neuron compare to that of H/R vehicle [(−) ST no H/R)] that is concomitant to previous finding of antioxidant potentials of ST [7, 56]. In addition, for better understanding how ST protects ROS production after H/R we further analyzed the expression of PINK1 as ROS are promising candidates of the physiological triggers for robust increase of mitophagy by PINK1dependent Parkin pathway [57, 58], with ubiquitin-binding adaptor p62 recruits cargo into autophagosome by binding to LC3; finally, degrade the mitochondria by lysosome [59]. In the present study we found also ST significantly induces the autophagy marker protein LC3II as well as PINK1 and their clearance and further investigation it makes clear that this autophagy induction was not chaperone dependent one i.e. HSC70 but through PINK1 dependent mechanism which is familiar to the NR2B-containing NMDA receptors mediated control of PINK1 expression in OGD-induced ischemic model [60]. Furthermore, apoptosis involving mitochondria is one of the cascade events that follow metabolic change during stroke activates proteolysis caspase-dependent DNase, resulting in internucleosomal fragmentation of DNA [61]. In this study we also found that ST treatment suppresses the expression of H2Ax punta expression after H/R compare to vehicle [(−) ST H/R)] indicating STs role in protection of DNA fragmentation.

It is well known that liver X receptors (LXRs) protects against cerebral ischemia mediated brain damage [62–64] and protect against glutamate toxicity by selective inhibition of Glun2B and induces mitophagy [47]. Since our previous transcriptomics [15] and report of Sabeva *et al*., [46] it has appeared that ST differentially influence ABC transporter expression therefore it might work as LXRβ agonist and helps in recovering H/R. From our molecular dynamics simulation observation, we found ST formed similar binding orientation to the known agonist desmosterol to LXRβ (Figure 6a). More precisely, guided from docking simulation, revealed that ST formed high number of hydrogen bonding to the GLU281 and ARG319 residues (Figure 6a), which indicates the similar orientation to the other endogenous steroids-nuclear receptor complexes [65]. Although docking study showed the hydroxyl group in position C-3 of the steroid skeleton made hydrogen bonding with the ASN239 and PHE329 residue, this group made strong hydrogen bonds with GLU281 on helix 3 and ARG319 on helix 5 of LXRβ on 100 ns long molecular dynamics simulation (Figure 1d)[66].

Excitotoxicity during an acute episode of stroke causes rapid elevation of glutamate levels as well as subsequent production of massive amount of ROS in the ischemic region of the brain. Thus inhibition of ischemic glutamate release could confer neuroprotection by terminating multiple downstream death signaling cascades at their converging upstream initiation point. Our present data shows that after H/R, ST can decreases neuronal toxicity and their subsequent exacerbating effect to cellular damages by protecting mitochondria, therefore could be a potential therapeutic for ischemic stroke.

## Supplementary document

**Table S1:** Result of glide docking analysis for stigmasterol and reference cocrystal agonist.

| Compound Name | Docking Score | Ligand efficiency | XP GScore | GScore | Glide energy | Glide emodel |
| --- | --- | --- | --- | --- | --- | --- |
| Ref. | -12.67 | -0.43 | -12.67 | -12.67 | -47.89 | -77.19 |
| Stigmasterol | -13.33 | -0.44 | -13.33 | -13.33 | -51.66 | -84.28 |

**Table S2:**
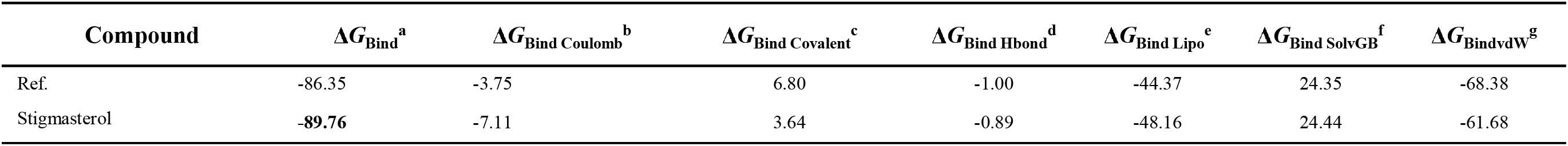
Relative binding free analysis by MM-GBSA

